# An analysis of respiratory function and mitochondrial morphology in *Candida albicans*

**DOI:** 10.1101/697516

**Authors:** Lucian Duvenage, Daniel R. Pentland, Carol A. Munro, Campbell W. Gourlay

## Abstract

Respiratory function and mitochondrial dynamics have been well characterised in a number of cell types, including the model yeast *Saccharomyces cerevisiae*, but remain under-researched in fungal pathogens such as *Candida albicans.* An understanding of mitochondrial activity and morphology is important if we are to understand the role that this organelle plays in adaption and response to stress. Here we examine the respiratory profiles of several prominent pathogenic *Candida* species and present a useful GFP probe for the study of mitochondrial morphology. We examine mitochondrial morphology under a variety of conditions that *Candida* species may encounter within the host, such as acidic pH, respiratory and oxidative stress. The GFP probe also allowed for the visualisation of mitochondria during hyphal development, during growth following macrophage engulfment and distribution within biofilms. These data demonstrate that the mitochondrial network of *C. albicans* is highly responsive to both environmental conditions and developmental cues, suggesting important roles for this organelle in environmental adaption.

## Introduction

A number of species of *Candida* species are known fungal pathogens of humans and are capable of causing infections that can be superficial, recurrent or indeed life-threatening. While *C. albicans* is the most frequently isolated species in cases of bloodstream infections there has a been an increase in the incidence of non-*albicans* infections (Montagna *et al.* 2014). An important aspect of Organisms that adopt a *C. albicans* is a Crabtree Negative yeast that relies on respiration to supply energy for proliferation and morphogenesis. A better understanding mitochondrial biology could open the possibility of new antifungal strategies against *Candida* infections.

The respiratory chain of *C. albicans* is able to adapt rapidly upon inhibition of certain components of the chain in order to allow respiration to continue. *C. albicans* possesses a classical electron-transfer chain (ETC) conserved in most eukaryotes but also an alternative pathway which allows is induced when the classical ETC is inhibited (Huh and Kang 2001). *C. parapsilosis* has been shown to possess a branched respiratory chain as well, with three pathways: classical, alternative and parallel. The contribution of the parallel pathway is only effective in situations where the other pathways are blocked, nevertheless it allows respiration to continue (Milani *et al.* 2001). Studies that have found the absence of complete inhibition in the presence of cyanide and alternative oxidase inhibitors suggest that a similar parallel pathway exists in *C. albicans* (Cavalheiro *et al.* 2004; Helmerhorst *et al.* 2005). The discovery of a regulator of Complex I as well as subunits of Complex I which are fungal specific raises the possibility of targeting respiration as an antifungal therapy (Calderone, Li and Traven 2015).

Mitochondria are dynamic organelles that constantly undergo fission and fusion. In yeast, fission of the mitochondrial network is an active process and a general response to stress (Knorre *et al.* 2013). Fission has been shown to isolate defective parts of the network and facilitate their turnover by mitophagy (Mao and Klionsky 2013). In recent years, much of the machinery governing fission and fusion has been elucidated (Pagliuso, Cossart and Stavru 2018), including important contributors such as interaction with the ER (Tucey *et al.* 2016). As mitochondrial function is implicated in a wide variety of pathologies and stress responses, the study of mitochondrial morphology in pathogens such as *C. albicans* could be used to better understand the host-pathogen interaction. For example, mitochondrial fragmentation in *Aspergillus fumigatus* has recently been used as a marker to study its interaction with human granulocytes (Ruf, Brantl and Wagener 2018).

Dyes such as rhodamine and MitoTracker have been widely used to enable visualisation of mitochondria. However, some dyes are dependent on the existence of mitochondrial membrane potential and therefore may not be suitable in certain studies of cells treated with inhibitors that collapse this membrane potential. Using a recombinant GFP system which targets GFP to mitochondria can be integrated into the genome of yeasts such as *C. albicans*, allowing the study of mitochondrial morphology under any conditions without the need for staining optimisation. Several GFP fusion proteins in *C. albicans* have shown mitochondrial localisation, including Tom70 (Sun *et al.* 2017), Srr1 and (Mavrianos *et al.* 2013) Goa1(Bambach *et al.* 2009). The fusion of GFP to a protein could also negatively affect function, or lead to changes in stability and expression level. The mitochondrial targeting of GFP is therefore more desirable. Mitochondrial probes have been used to study the effects of environmental stresses and treatments in other systems. These include the effects of toxins (McLaughlin *et al.* 2009) and oxidative stress (Dong *et al.* 2015).

In this study we observe differences in the respiration profiles of *Candida* species, and the adaptability of respiration in *C. albicans* to extreme low pH conditions. We used a GFP probe to examine mitochondrial morphology in *C. albicans* under stress conditions, including low pH, respiratory- and oxidative stress and observed fragmentation of the mitochondrial network in common with these responses. We also examined mitochondrial dynamics during the yeast-to-hypha transition and mitochondrial distribution in biofilms. Overall, our study of the adaptability of respiration behaviour and mitochondrial dynamics enhances our appreciation of role of mitochondria in stress responses in a fungal pathogen, which could be exploited in development of new antifungal therapies.

## Materials and Methods

### *Candida* strains, growth and chemicals

The *C. albicans* strains SC5314 and BWP17 and *C. glabrata* BG2 were used in this study. The *C. tropicalis, C. parapsilosis* and *C. guillermondii* strains used are clinical isolates, a kind gift from Prof. Judith Berman (Department of Molecular Microbiology and Biotechnology, Tel Aviv University). *Candida* strains were maintained on YPD agar plates and grown in YPD in a 30 °C shaking incubator unless stated otherwise. The concentrations of sodium nitroprusside dihydrate (SNP) and salicylhydroxamic acid (SHAM) (Cat. No. 1.06541 and S607, Sigma-Aldrich, Dorset, UK) used for inhibition were 1 mM and 0.5 mM respectively. 2-Thenoyltrifluoroacetone (TTFA) (Cat. No. T27006), Triethyltin bromide (TET) (Cat. No. 288047), FCCP (carbonylcyanide p-trifluoromethoxyphenylhydrazone) (Cat. No. C2920) Potassium cyanide (KCN) and hydrogen peroxide were obtained from Sigma-Aldrich.

### Whole cell respirometry

Respiromety was carried out in real time using an Oxygraph-2k respirometer (Oroboros Instruments, Austria) which was calibrated at 30 °C as per the manufacturer’s instructions. Cells from an overnight culture in YPD were added to 3 ml fresh YPD to a final optical density at 600 nm (OD_600_) of 0.2. The cells were incubated at 30 °C with shaking for 2 hours. The cells were then counted and diluted in YPD to give a final cell concentration of 1 × 10^6^ cells/ml, of which 2.5 ml was added to each chamber in the respirometer. The respiration was allowed to reach routine levels before the addition respiration-modulating drugs as follows: 160 µM triethyltin bromide (TET), 10 µM FCCP, 1 mM KCN, 1 mM SHAM. Data was analysed using Datlab 6 software (Oroboros Instruments). Three independent experiments were performed per species. To examine the effect of low pH on respiration, cells were incubated in YPD buffered with 50 mM MOPS (pH 6) or 200 mM glycine-HCl (pH 2) for 18 h at 30 °C. Cells were then transferred to fresh media of the same respective types and incubated at 30 °C for 2 h prior to respirometry. Four independent experiments were performed per condition. Student’s t test was used to compare groups.

### Construction of mitochondrial GFP strain

In order to examine the effects of respiration inhibition and hyphal growth on mitochondrial morphology in *C. albicans*, GFP was targeted to the mitochondrial matrix. To achieve this, the predicted signal sequence from *C. albicans ATP5* (MISRVFSRSLASAAKSTKPPIQLFGID) was added upstream of the GFP sequence in the plasmid pACT1-GFP, a kind gift from Professor A. Brown (Barelle *et al.* 2004). The signal sequence was predicted using MitoProt II v1.01 (Claros and Vincens 1996) and the coding sequence, including HindIII restriction sites and alanine linker sequence was synthesized by GeneArt (Thermo Fisher Scientific, UK): AAGCTTTATTAAAATGATTTCTCGTGTTTTTTCCCGTTCATTAGCATCAGCTGCCAAGTCTACCAAACCACCAATCCAATTGTTTGGTATTGATGCAGCTGCAGCTGCAGCAAGCTT. This sequence was digested with HindIII and cloned into pACT-1 GFP. The resulting construct was named pACT1-mtGFP. A schematic of the expression construct is shown in Figure 2B. pACT1-mtGFP-mtGFP was linearised by digestion with StuI for integration at the RPS10 locus as described in (Barelle *et al.* 2004) and transformed into BWP17 using an electroporation protocol (Thompson *et al.* 1998). Transformants were selected using agar plates containing Yeast Nitrogen Base without amino acids supplemented with 2% glucose and -URA dropout powder (Formedium, UK). The resulting strain BWP17-mtGFP was used for all mitochondrial morphology studies.

**Figure 1.**
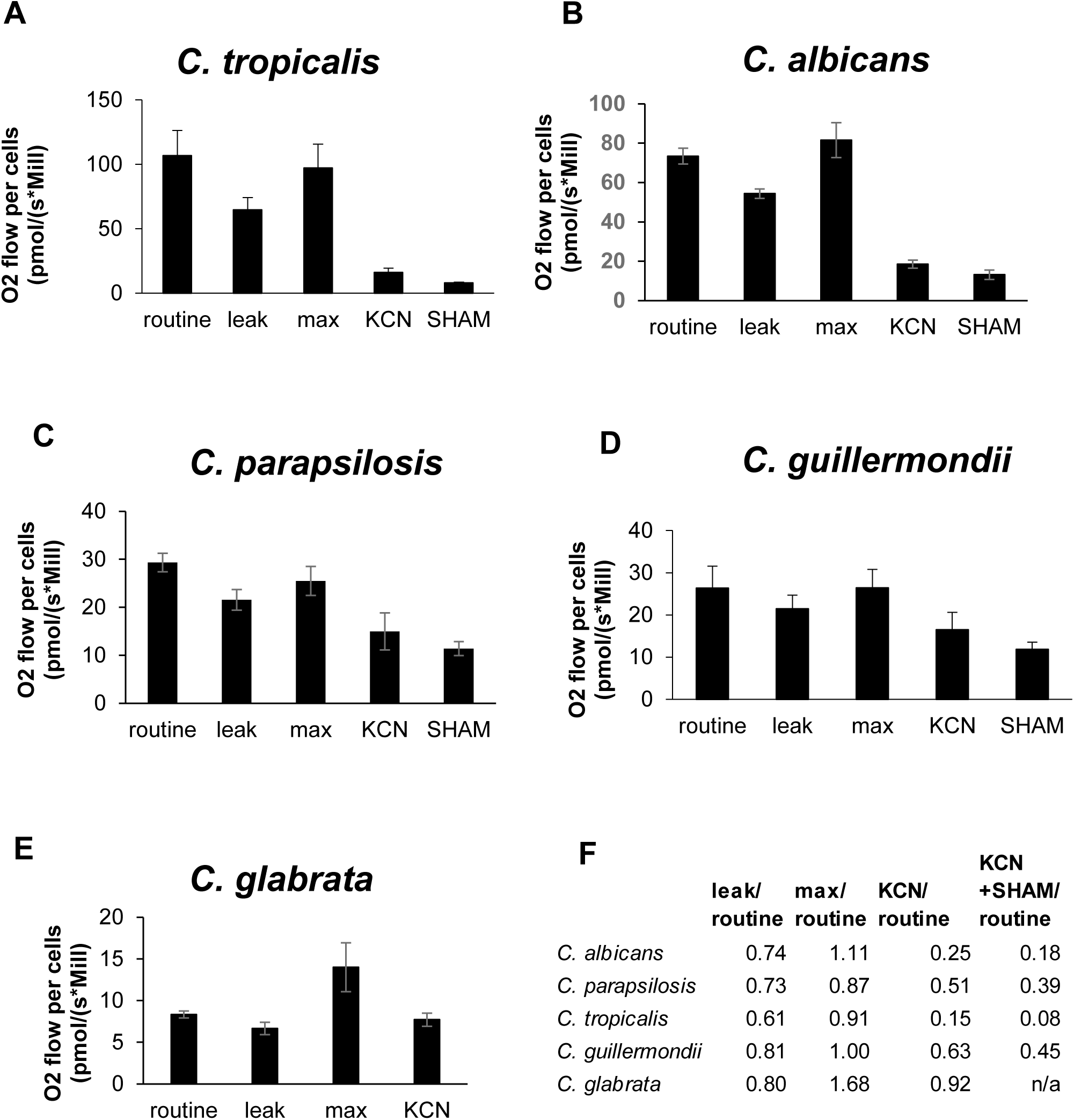
Whole-cell respirometry in *Candida* species. Routine respiration was recorded immediately before the addition of drugs, “leak” respiration was recorded after addition of ATP synthase inhibitor triethyltin, and “max” respiration represents maximum electron transport chain capacity in the presence of uncoupler FCCP. Respirometry was performed for each *Candida* species (**A-E**) using the drug schedule described in Materials and Methods. **F.** Summary of ratios of inhibited or uncoupled mitochondria to routine respiration for each species. Three independent experiments were performed per group.

**Figure 2.**
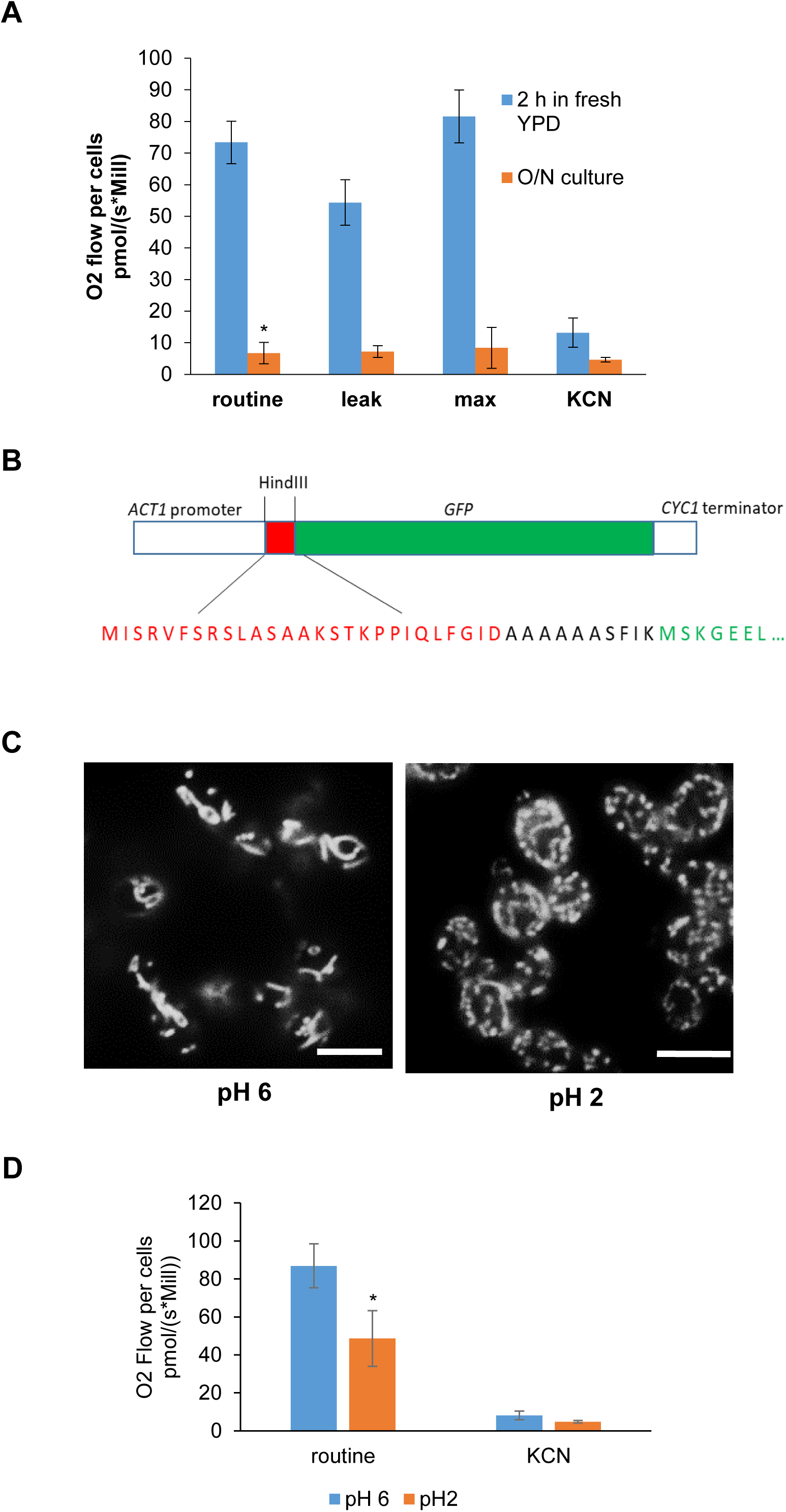
Respiration and mitochondrial morphology at low pH. **A.**Whole cell respirometry of *C. albicans* comparing respiration of cells from an overnight culture in YPD to those recovered in fresh YPD for 2 h, n=3, *p<0.01. **B.** Schematic of the mitochondrial targeted GFP construct in pACT1-mtGFP, showing *C. albicans ACT1* promoter and *CYC1* terminator sequences, predicted mitochondrial targeting sequence in red and GFP in green. **C.** Representative examples of mitochondrial morphology in pH 2 and pH 6 cells. Bar = 10 µm. **D.** Respirometry of *C. albicans* incubated in buffered YPD at pH 2 or pH 6, showing routine- and 1 mM KCN-inhibited respiration rates, n=4, *p<0.05.

### CFU viability assays

Cells from an *C. albicans* SC5314 overnight culture in YPD were diluted in YPD to a final OD_600_ of 0.2. The cells were grown for 5 hours at 30 °C with shaking. KCN and SHAM were added to a final concentration of 1 mM and 0.5 mM respectively and cultures were grown for a further 18 h. The cells were collected, washed three times in PBS and counted using a haemocytometer, then diluted in PBS. Four hundred cells of each suspension were plated on YPD plates. The plates were incubated for 24 h and the number of colony forming units (CFU’s) were counted. Three independent experiments were performed. Student’s t-test was used to compare groups.

### Hyphal growth and chemical treatments

Examination of mitochondrial morphology at low pH was performed by incubating cells in YPD buffered with 50 mM MOPS (pH 6) or 200 mM glycine-HCl (pH 2) for 18 h at 30 °C. Cells were washed three times in PBS and examined by microscopy using the GFP filter on an Olympus IX81 inverted microscope, illuminated using a CoolLED pE-4000 unit and captured using an ANDOR Zyla 4.2 CMOS camera. To examine mitochondrial morphology during the course of hyphal growth, cells from an overnight culture in YPD were washed three times in PBS and used to inoculate RPMI-1640 media (Sigma-Aldrich, R8755) to a final OD_600_ of 0.2. Cultures were grown at 37 °C in a shaking incubator. Samples were taken at regular intervals and examined by microscopy. To examine the effect of respiratory stress on mitochondrial morphology in yeast cells, log phase cells in YPD were treated with 1 mM SNP, 0.5 mM SHAM or 50 µg/ml TTFA for 18 h. The cells were washed three times in PBS and examined by microscopy. To examine the effects of peroxide, cells from an overnight culture in YPD were washed three times in PBS and used to inoculate synthetic complete medium, 2% glucose, followed by incubation at 30 °C for 5 hours. Cells were then treated with 5 mM hydrogen peroxide for 2 hours followed by microscopy. The procedure used for hyphal cells was the same except that RPMI and an incubation temperature of 37 °C were used.

### Phagocytosis by of *C. albicans* by murine macrophages

J774.1 murine macrophages were maintained in DMEM with 10% FBS, 200 U/ml penicillin/streptomycin (respectively, Cat No. 10569010, 10082147, 15070063, Gibco, Thermo Fisher) at 37 °C, 5% CO_2_. Cells were counted and diluted in fresh medium to give 5 × 10^4^ cells in 0.3 ml which was added to the wells of a µ-Slide 8 Well (ibidi GmbH, Germany). The cells were then incubated overnight. *C. albicans* BWP-17 mtGFP cells from an overnight culture were washed three times in PBS and counted, then diluted to 1.5 × 10^4^ cells in 0.3 ml in macrophage culture media and vortexed briefly. This cell suspension was then added to macrophages. The cells were then co-incubated for 3 h at 37 °C, 5% CO_2_ and examined by microscopy using a GFP filter and low level brightfield illumination.

### Biofilm growth and imaging

Biofilms were grown on microscope slides coated with polydimethylsiloxane (PDMS) silicone in petri dishes: The surface was immersed in 50 % FBS in PBS for 30 min at 30 °C, then washed twice with PBS. BWP17-mtGFP cells from an overnight YPD culture were washed three times in PBS, adjusted to an OD_600_ of 1.0, and allowed to adhere to the surface for 90 min at 37 °C. Non-adherent cells were removed by washing with twice with PBS. Adhered cells were immersed in RPMI-1640, followed by incubation at 37 °C for 48 h. The biofilm was washed twice with PBS, then stained with of 50 µg/ml ConA-Texas Red (Thermo Fisher, Cat no. C825) in PBS, with 45 min incubation in the dark at room temperature. The biofilm was washed twice with PBS and samples were mounted: Two drops Prolong Diamond antifade mountant (Thermo Fisher Scientific, Cat. No. P3696) were added to the biofilm, a coverslip was added and the samples were left to cure overnight in the dark at room temperature. Biofilms were imaged using a Zeiss LSM880/Elyra/Axio Observer.Z1 confocal microscope with the 488nm (GFP) and 561nm (Texas-Red) laser lines. Z-stack images were obtained at intervals of 1.4 µm over a range of 30 µm. Z-stack reconstruction was performed using ZEN software (Blue edition) (Zeiss, Germany).

## Results and discussion

### Characterisation of respiration in *Candida* species

We wished to characterise respiratory chain function in a number of pathogenic *Candida* species. Triethyltin Bromide (TET) was used to inhibit ATP synthase, allowing measurement of oxygen flux under conditions of inhibited proton translocation by ATP synthase. Following TET addition protons may enter the matrix only by diffusion (“leak” respiration) giving an assessment of how well electron transport is coupled to ATP production. FCCP is a proton-ionophore and was used to measure uncoupled respiration, or maximal capacity of the electron transport chain (“max”). The classical electron transport chain (ETC) was inhibited at the level of Cytochrome c Oxidase using KCN, and the alternative oxidase inhibitor salicylhydroxamic acid (SHAM) was used to inhibit remaining respiration contributed by the alternative pathway.

Among the *Candida* species measured, *C. tropicalis* exhibited the highest routine respiration rate (expressed as O_2_ flow per 10^6^ cells (pmol/(s*Mill)) when grown in rich YPD media, 107.6 ± 19.7 (Figure 1A), followed by *C. albicans*, 73.4 ± 4.0 (Figure 1B). Interestingly *C. parapsilosis* and *C. guillermondii* exhibited significantly lower routine respiration levels, 29.3 ± 1.9 and 26.3 ± 5.2 respectively (Figure 1C and 1D). *C. glabrata* exhibited very low routine respiration, (8.3 ± 0.4) most likely attributable to its preference for aerobic glycolysis (Figure 1E). All strains responded to TET inhibition and FCCP uncoupling of respiration, showing that the respiratory chain was intact and functional. It is worth noting that *C. tropicalis* exhibited the lowest routine/leak respiration ratio indicating that it exhibited the most efficient coupling of electron transport to ATP production (Figure 1A). We also observed that all Candida strains tested, with the exception of *C. glabrata*, had max/routine ratios that were close to 1 (Figure 1F). A max/routine ration of close to 1 suggests that electron transport is occurring at or close to its maximum in *C. albicans, C. tropicalis, C parapsilosis* and *C. guillermondii* during growth in YPD. In contrast the higher ratio observed in *C. glabrata* is likely to be indicative of a repressed ETC. The electron transport profile of C. glabrata is very similar to that of the non-pathogenic yeast *S. cerevisiae*, which also prefers to utilise fermentation during growth. All species tested responded to KCN inhibition, although the degrees of sensitivities were different. For example, *C. parapsilosis* and *C. guillermondii* were more resistant to KCN than *C. albicans*, as shown by the KCN-treated/routine respiration ratios (Figure 1F). The alternative and parallel respiratory pathways which have previously been characterised in *C. parapsilosis* (Milani *et al.* 2001) may be more active than in *C. albicans* under these conditions. The alternative oxidase inhibitor SHAM further decreased respiration following KCN addition, including in *C. guillermondii* and *C. tropicalis*, suggesting that the alternative oxidase is conserved in these species as well. Interestingly *C. glabrata* also exhibited resistance to KCN and suggests that the majority of oxygen consumption during growth in YPD does not occur via the mitochondrial Cytochrome c oxidase (Figure 1E).

The relatively high level of respiration in *Candida* species with glucose as a carbon source, at least when compared to Crabtree-positive organisms such as *S. cerevisiae*, suggest that respiration plays a major role in energy generation for these organisms under the conditions tested, with the exception of *C. glabrata*, which has also been shown to repress respiration in high glucose (Legrand *et al.* 2016). A number of Candida species, such as *C. albicans*, appear to rely on mitochondrial respiration for normal growth. This suggests that mitochondria may provide a novel target for therapy. However it should be noted that these data do not support a strict correlation between respiratory profile and pathogenicity.

### Mitochondrial function and morphology in *C. albicans*

Next we focussed on examining respiratory function and mitochondrial morphology in more detail in the most common yeast pathogen of humans, *C. albicans*. Initial experiments with *C. albicans* showed that cells grown overnight to stationary phase culture exhibited negligible levels of oxygen consumption that were attributable to mitochondrial electron transport (Figure 2A). As there was no effect on oxygen consumption when TET or FCCP was added to stationary phase *C. albicans* cells it can be stated that mitochondrial ETC activity does not support ATP production in stationary phase. The suppression of ETC was rapidly reversed upon sub-culture into fresh media and within 2 h mitochondrial electron transport was operating at its maximum rate (Figure 2A). This significant change highlights that mitochondria are activated upon stimulation of growth, but also that care that must be taken when investigating respiratory levels in living cells. Our data also supports previous observations that respiration declines in stationary phase *C. albicans* cells (Helmerhorst, Troxler and Oppenheim 2001).

Mitochondrial dynamics are an important indicator of cellular quality control (Westermann 2010). The degree of fission of the mitochondrial network can provide information on the metabolic status of the cell. In yeast, mitochondrial fission is a general stress response which attempts to isolate defective mitochondria for autophagy (Knorre *et al.* 2013; Mao and Klionsky 2013). We constructed a GFP probe to label the mitochondria in order to study the changes in mitochondrial morphology in various stress conditions and during the yeast-to-hypha transition in *C. albicans.* This was achieved by of the fusion of the N-terminal signal peptide for Atp5, a subunit of the ATP synthase, to GFP for targeting to the mitochondrial matrix. A schematic of the expression construct is shown in Figure 2B. Mitochondrial GFP probes have been shown to be useful tools in *S. cerevisae* and in mammalian cells. To target GFP to the mitochondria in *S. cerevisiase*, first 69 amino acids of the *Neurospora crassa* F_0_ ATP synthase subunit 9 was fused to the N-terminus of GFP (Westermann and Neupert 2000). The successful use of this well-characterised mitochondrial targeting sequence in a different species shows that mitochondrial targeting in fungi is relatively well-conserved. To target GFP to mitochondria in mammalian cells, a portion of the *COX8* precursor sequence was used (Rizzuto *et al.* 1995).

*C. albicans* encounters a variety of different environments as a pathogen, including fluctuations in pH, from pH 2 in the stomach (Zwolinska-Wcisło *et al.* 2001) to pH 6 in the oral mucosa (Cannon and Chaffin 1999). To investigate the impaired respiration observed in conditions of extreme low pH, mitochondrial morphology was examined in cells exposed to a pH of 2. These cells showed a more fragmented mitochondrial network than those grown at pH 6 (Figure 2C). This may suggest that low pH can induce mitochondrial damage or indeed that it initiates a programme of fission as part of a response to this stress. To examine the effect of extreme low pH on respiration rate, cells were grown at pH 2 in buffered YPD for 18 h at 30 °C, followed by growth at pH 2 in fresh media for 2 hours prior to respirometry. Respirometry showed that routine respiration was reduced by approximately 50 % in pH 2 cells (Figure 2D). These results show that extreme low pH can affect mitochondrial function. Respiration could be inhibited by KCN under all conditions of growth suggesting that *C. albicans* uses ETC function over range of pH values.

### Mitochondrial morphology in *C. albicans* hyphae

The inhibition of respiration in *C. albicans* has been shown to prevent filamentation (Grahl *et al.* 2015) and measurement of respiration rates in hyphae show an increase in oxygen consumption relative to yeast cells (Guedouari *et al.* 2014). Respiration and mitochondrial activity are therefore essential for the switch from yeast to hyphae. In order to examine the changes in mitochondrial morphology that occur during the yeast-to-hypha transition, cells were grown in RPMI at 37 °C and were imaged at regular intervals. The mitochondrial network was seen to extend into new germ tubes as soon as they were formed (Figure 3A). The characteristic mitochondrial network in yeast cells is dense and continuous and this was observed within hyphal cells as they extended for up to 2h. However, after 6h of growth in hyphal induction conditions, the mitochondrial network, although still continuous and fused, appeared notably less dense (Figure 3A). These data support a model whereby mitochondrial biogenesis is restricted to the growing end of hyphal cells as they extend.

**Figure 3.**
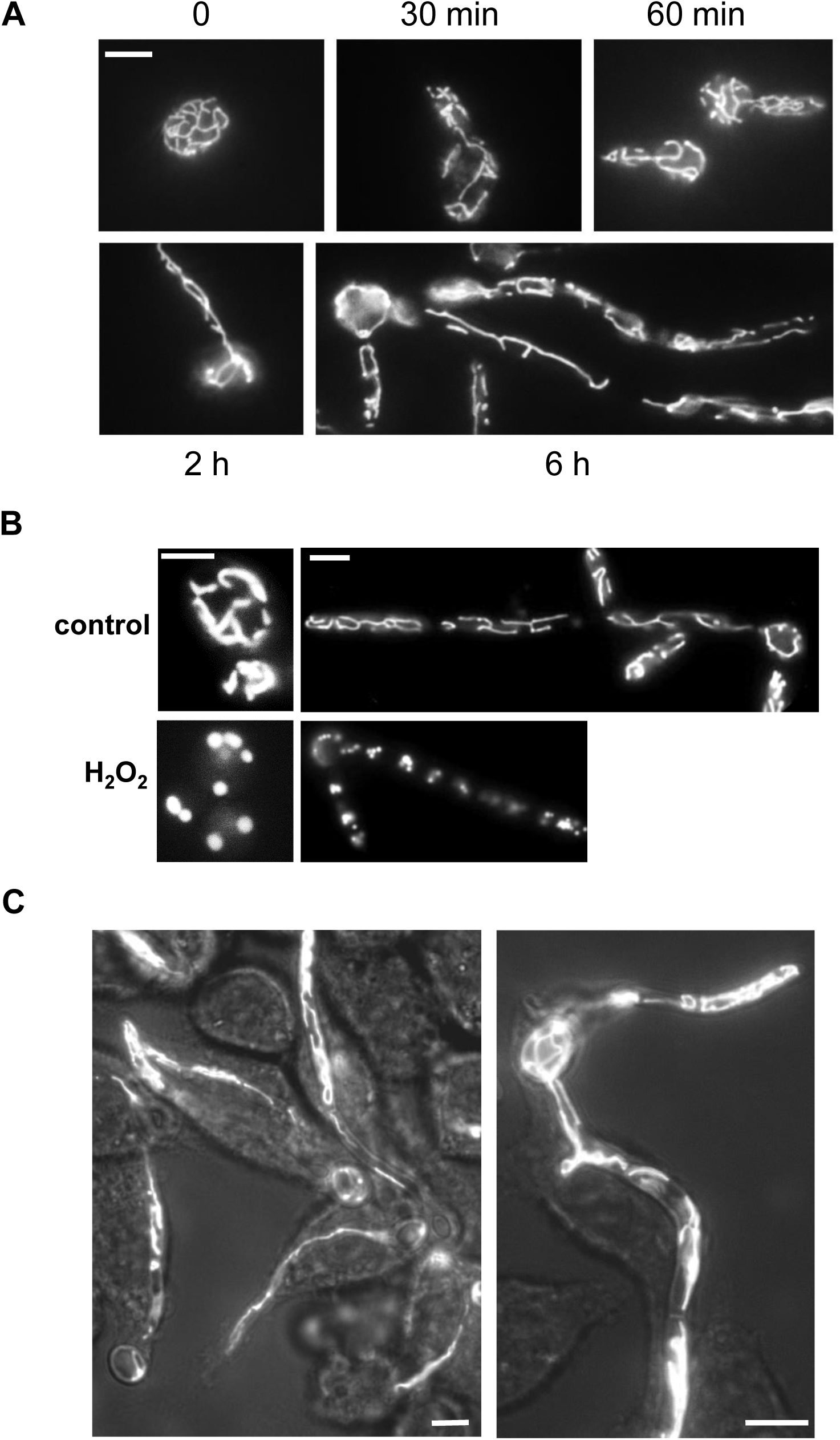
*C. albicans* mitochondrial morphology during yeast-to-hypha transition and in hyphae under stress. **A.** Time course of *C. albicans* filamentation in RPMI at 37 °C examining mitochondrial morphology. **B.** Comparison of mitochondrial morphology in yeast and hyphae treated with 5 mM hydrogen peroxide for 2 h. Bar = 10 µm. **C.** Representative examples of *C. albicans* hyphal growth inside J774 macrophages following 3 h co-incubation. Bar = 10 µm.

As was the case for yeast cells exposed to low pH stress we found that the mitochondrial network altered its morphology in response to a range of stress in hyphal cells. Both yeast and hyphae exposed to hydrogen peroxide showed a fragmented network of clustered mitochondria (Figure 3B) which was also observed with hydrogen peroxide treatment of *S. cerevisiae* (Karatani *et al.* 2013) and *A. fumigatus* (Ruf, Brantl and Wagener 2018). Study of the oxidative stress response in *S. cerevisiae* suggests that this fragmentation is dependent on nuclear-to-cytoplasmic translocation of the transcriptional repressor cyclin C, which as a downstream effector of the mitogen-activated protein kinase Slt2 is integrated with the cell wall integrity response (Jin, Strich and Cooper 2014).

*C. albicans* hyphae growing in macrophages did not show an altered mitochondrial morphology compared to hyphae grown in RPMI (Figure 3C), suggesting that these cells were defended against oxidative stress, or that the oxidative stress within the phagosome was not high enough to affect mitochondrial morphology. The J774A macrophage line has been described as having a weak oxidative burst; they are unable to kill *C. albicans* (Lorenz, Bender and Fink 2004). A similar defect caused by hydrogen peroxide treatment was observed in *ccz1*Δ *C. albicans* mutant but not in the corresponding in wild type, which the authors ascribed to a loss of membrane potential (Dong *et al.* 2015). The difference in response of wild-type *C. albicans* to H_2_O_2_ in this study could have been due the use of synthetic media instead of YPD. Fragmentation of the mitochondrial network in response to the oxidative agent *tert*-butyl hydroperoxide was described in another dimporhic fungus, *Y. lipolytica* (Rogov *et al.* 2016). *C. albicans* treated with diorcinol D also exhibited an altered mitochondrial morphology as shown by the distribution of a Tom70-GFP fusion which was attributed to an increase in oxidative stress (Li *et al.* 2015). *C. albicans* has been shown to delay phagosome maturation, including acidification, during hyphal growth inside macrophages (Bain *et al.* 2014). This could explain why the fragmentation of the mitochondrial network that we observed at low pH did not occur.

Fragmentation of the mitochondrial network was also observed when cells were treated with respiration inhibitor SNP (which generates Nitric Oxide), the alternative oxidase inhibitor SHAM or the complex II inhibitor TTFA (Figure 4A-C), suggesting that fragmentation can also result from impaired respiration at multiple points within the respiratory system. These responses may be physiologically relevant, as macrophages impose nitrosative stress on *C. albicans* in the phagosome which can disrupt the respiratory chain. Growth for 18 h in the presence of inhibitors to block both ETC and alternative oxidase function (SNP+SHAM) also led a proportion of cells showing a loss of mitochondrial structural integrity that may be indicative of cell death (Figure 4C). GFP signal not associated with cells was also detected, indicating a degree of cell lysis and release of mitochondrial components (Figure 4C). These data suggest that prolonged inhibition of mitochondrial respiration can lead to cell death in *C. albicans*, In support of this we observed that treatment of *C. albicans* with KCN, to which they are resistant as a result of alternative oxidase upregulation, or KCN + SHAM (Aox inhibitor) for a prolonged period led to a significant loss of viability (Figure 4D).

**Figure 4.**
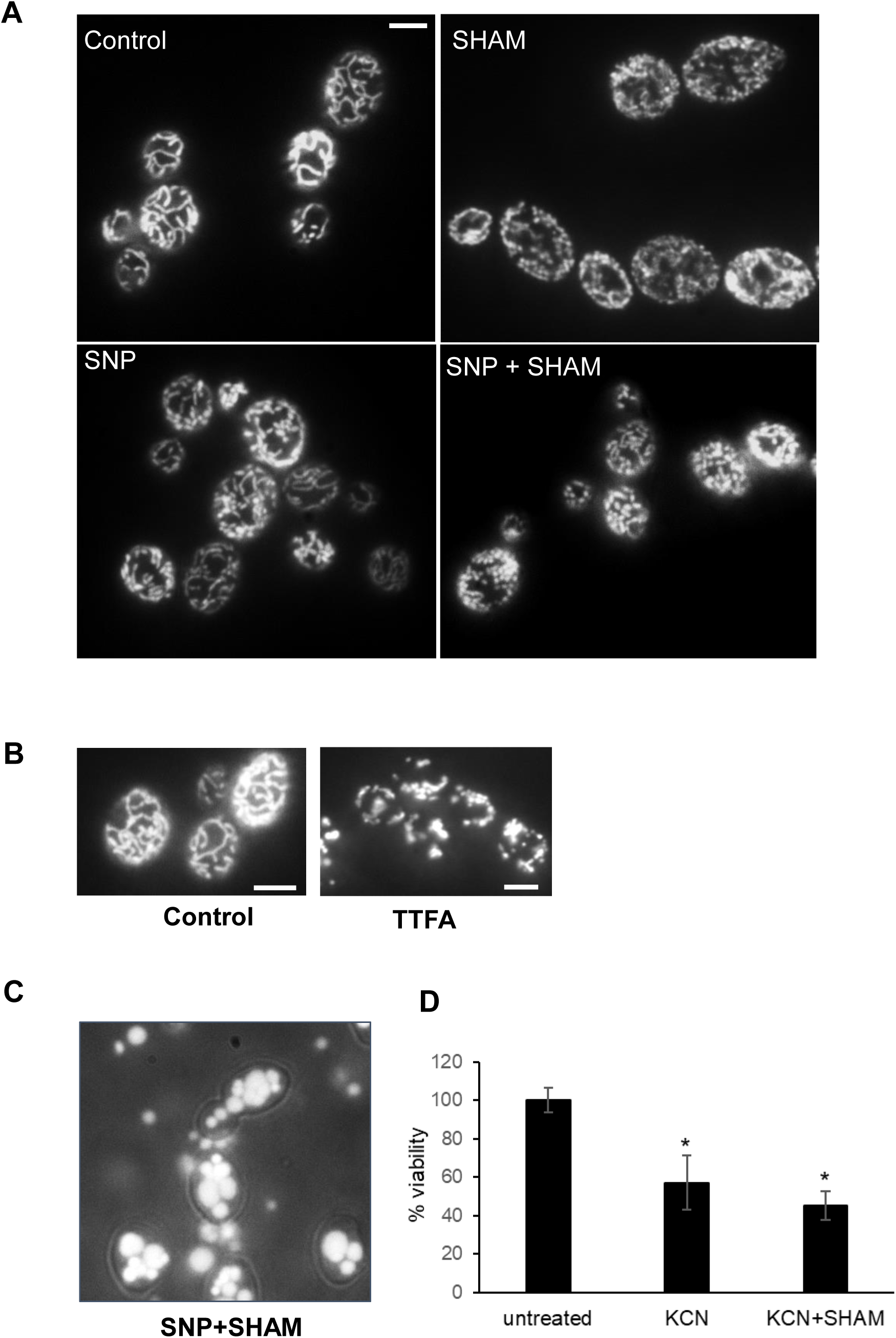
Mitochondrial morphology in respiratory stress. Treatment of *C. albicans* with respiration inhibitors: **A.** 1 mM SNP, 0.5 mM SHAM, or both for 18 h and examination of mitochondrial morphology. **B.** Treatment of *C. albicans* with 50 µg/ml TTFA (Complex II inhibitor). **C.** Representative example of loss mitochondrial structure and cell lysis observed in a proportion SNP+SHAM treated cells. **D.** Viability determined by CFU counts after treatment with 1 mM KCN and in combination with 0.5 mM SHAM for 18 h, n=3, *p<0.01. Bar = 10 µm.

### Mitochondrial distribution within *C. albicans* biofilms

Biofilm formation by *Candidia* species on abiotic surfaces contributes to failure of medical devices such as voice prostheses (Talpaert *et al.* 2015), and indwelling catheters are a major source of invasive *Candida* infections (Kojic and Darouiche 2004). We used the BWP17-mtGFP strain to examine mitochondrial morphology in mature biofilms formed on a silicone surface. Mitochondria appeared fragmented and clustered (Figure 5A, B), a phenotype similar to our observation of peroxide-treated hyphae (Figure 3B). Given the fused appearance of mitochondria in planktonic hyphal cells (Figure 3A), it is possible that mitochondrial morphology is significantly altered in within biofilms. However, it is also possible that our sample mounting procedure imposes a stress that in turn leads to fragmentation of the network. Mannan staining with fluorescent ConA showed that hyphae were more abundant towards the top section of the biofilm (Figure 5C), consistent with published observations. Interestingly, although the mitochondrial GFP could be visualised throughout the biofilm, both in yeast and hyphal cells, the signal was more intense in the upper section, suggesting increased mitochondrial density at hyphal tips (Figures 5E, F). This could reflect increased mitochondrial biogenesis or respiratory activity within the upper section. This may be a consequence of its interaction with the oxygen-rich media, as compared to hypoxic environment which is thought to be generated in the interior of *Candida* biofilms (Rossignol *et al.* 2009).

**Figure 5.**
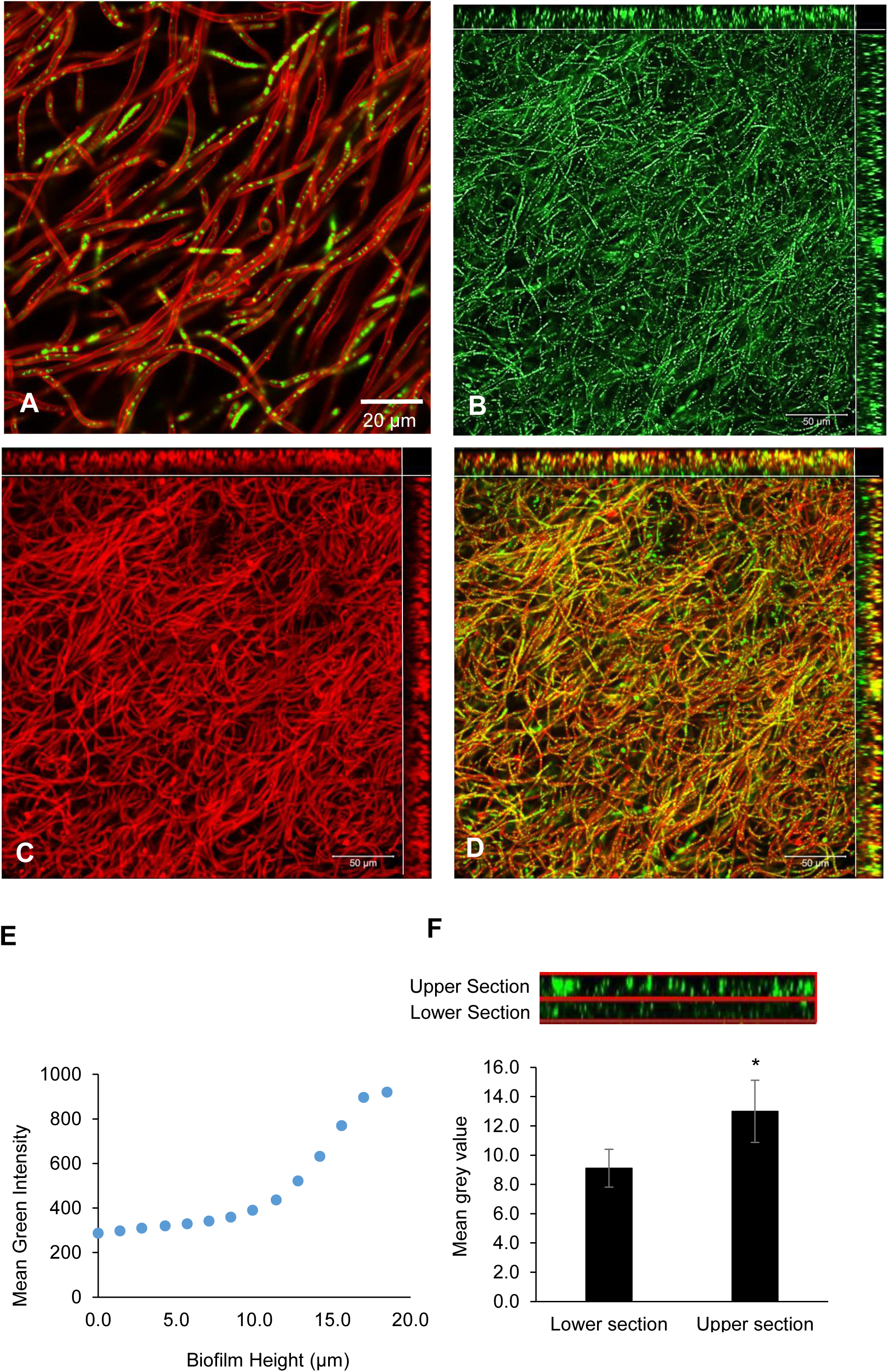
Mitochondrial distribuition in biofilms on a silicone surface. Biofilms of the BWP17-mtGFP strain formed on a silicone surface after 48 h were stained with ConA-Texas Red and examined by confocal microscopy. **A.** Mitochondrial distribution within biofilm hyphae. **B.** Distribution of mitochondria throughout the biofilm. The main panel is a top-down view of the biofilm and the top- and right-hand side panels depict the biofilm interior reconstructed from a z-stack. **C.** Distribution of hyphae throughout biofilm, shown by mannan staining. **D.** Merged hypha- and mitochondrial distribution. **E.** Representative analysis of fluorescence intensity (mean green value) measured for z-stack images with increasing biofilm height. **F.** Mitochondrial GFP fluorescence intensities (expressed as mean grey values) of the top and bottom halves of biofilm z-stack reconstructions were compared using ImageJ. Data represents means and standard deviations for three biofilm samples, *p<0.01.

## Conclusion

In this study we measured respiration in several pathogenic *Candida* species. We showed that most exhibited high levels of respiration that was linked to ATP production which is line with their Crabtree negative status. The exception to this was *C. glabrata*, which exhibited very low routine respiration, consistent with glucose repression and a lifestyle more reliant on sugar fermentation. As all species tested are human pathogens our data does not support a model whereby a specific respiratory strategy may act as a predictor of virulence. To visualise mitochondria under conditions of variable membrane potential we constructed a robust GFP probe to label mitochondria in *C. albicans.* This replaces the need for staining optimisation with dyes such as MitoTracker. We show that the mitochondrial network in *C. albicans* is continuous under normal growth conditions but becomes fragmented under conditions of stress. We observed that mitochondria within mature biofilms are distributed unevenly which may reflect oxygen availability within this structure. Further study of mitochondrial morphology and respiratory function under various stress conditions could provide insights into the roles of mitochondria in signalling and stress response that may facilitate future antifungal development and a better understanding of the cell biology of *Candida* species.

## Funding

This work was supported by a PhD studentship award to Lucian Duvenage within a Wellcome Trust Strategic Award for Medical Mycology and Fungal Immunology 097377/Z/11/Z.

## Acknowledgements

We would thank members of the Aberdeen Fungal Group, Kent Fungal Group and WTSA MMFI consortia for comments and useful discussion throughout the project.

